# Fung-AI: An AI/ML-driven pipeline for antifungal peptide discovery

**DOI:** 10.64898/2026.03.09.710548

**Authors:** Daniel S Berman, Libby M Lewis, Tom D Curtis, Olivia N Tiburzi, Daniel F Q Smith, Arturo Casadevall, Laura J Dunphy

**Affiliations:** Johns Hopkins Applied Physics Laboratory, 11100 Johns Hopkins Rd., Laurel, MD 20723, USA; W. Harry Feinstone Department of Molecular Microbiology and Immunology, Bloomberg School of Public Health, The Johns Hopkins University, Baltimore, Maryland, USA

**Keywords:** Generative Adversarial Networks, Sequence Generation, Antifungal Peptide, Deep Learning, Drug Discovery

## Abstract

Emerging fungal pathogens represent a concerning threat to both global health and food security. In this study, we aimed to address our rising vulnerability to fungal pathogens through the development of the Fung-AI pipeline: an AI/ML-driven approach for antifungal discovery. A generative adversarial network (GAN) was trained to generate novel candidate antifungal peptide sequences. Next, *in silico* antifungal and hemolytic classifiers were built to further prioritize AI-generated peptides for experimental validation. From a pool of ∼10,000 candidates, thirteen peptides were selected for testing over two-stages of experimentation. Five peptides were found to display mild antifungal activity against the wheat pathogen, *Fusarium graminearum*, with minimal inhibitory concentrations (MICs) ranging from 250 µg/mL to 500 µg/mL. Four of the five peptides also showed activity against the human pathogen, *Candida albicans* (MIC: 500 µg/mL). Two of our AI-generated antifungal peptides additionally demonstrated low cytotoxicity in HepG2 human liver carcinoma cells (LC50 > 704.2 µg/mL) indicating that they may be useful as scaffolds for future optimization for therapeutic applications. None of our peptides were found to considerably inhibit the emerging pathogen *C. auris*, suggesting the need for pathogen-specific down-selection of candidate peptides. Overall, we present a proof-of-principle, generative-AI-based approach for the rapid design of *de novo* antifungal peptides.

## Introduction

A major threat of growing global concern, fungal pathogens have been estimated to cause at least 2.5 million infection-related deaths per year (1–3), while additionally driving the spoilage of between 10-23% of annual pre-harvest crop yields (4). The negative impact of fungal pathogens has been exacerbated in recent years by both a general lack of known classes of antifungal drugs, as well as the emergence of multi- and pan-drug-resistant clinical isolates (5–8). Compounding upon this issue, the discovery of novel antifungal drugs is slow and challenging, in part because fungi share much of their cell biology with mammalian cells (9,10).

Antifungal peptides, short polypeptides that are typically 2-50 amino acids in length, are thought to be less likely to cause fungal resistance, and therefore represent a promising alternative to traditional small molecule drugs (11,12). Antifungal peptides, many of which are naturally produced by bacteria, plants, and animals, can kill fungal cells through a variety of mechanisms, including cell membrane disruption, lysis of the cell wall, and targeting of critical intracellular processes (12). Published examples have demonstrated strong activity against relevant human pathogens, including *Candida albicans, Aspergillus fumigatus*, and *Cryptococcus neoformans* (10). For example, the archaeal peptide, VLL-28 and a peptide isolated from the giant monkey frog, Skin-PYY, have been shown to have minimal inhibitory concentrations (MICs) of 88.5 µg/mL and 25 µg/mL against *C. albicans* (10,13,14), respectively. While some antifungal peptides can also exhibit high toxicity in mammals, this toxicity can sometimes be mitigated with the design of semi-synthetic or synthetic analogues (10,15).

Over the last decade, the rapid evolution of generative artificial intelligence (AI) has resulted in the explosion of AI-enabled research across the biological sciences. Generative modeling approaches, such as generative adversarial networks (GANs) (16), variational autoencoders (VAEs), and diffusion models (17), can be trained to turn random noise into realistic DNA, RNA, and protein sequences. For example, used in conjunction with other deep learning architectures, GANs have been applied to predict viral evolution (18), discover new enzymes (19), and optimize DNA and protein sequences for specific functions (20,21). More recently, deep learning has been used successfully to both detect (*e*.*g*., with discriminative classification models) (22–27) and design (*e*.*g*., with generative models) novel antimicrobial peptides (28–32). However, while there has been some success in developing highly accurate AI/ML models for the classification of antifungal peptides (33–39), the direct application of generative AI to antifungal discovery has been much more limited, at least in part due to a lack of large, high-quality fungal datasets (40).

Given the recent success of others toward developing high-accuracy antifungal peptide classifiers, we hypothesized that, while limited, there could be sufficient data in the public domain to train an antifungal peptide generative model for the task of drug discovery. To this end, we present the Fung-AI pipeline, an *in silico* approach for antifungal discovery (**Figure 1**). The Fung-AI pipeline begins with a GAN, which has been trained on publicly available known antifungal and non-antifungal peptide sequences to rapidly generate diverse and novel candidate peptide sequences. Peptides are then down-selected based on predicted activity, toxicity, structure, as well as computed biophysical properties (*e*.*g*., net charge, hydrophobicity). To validate our approach, thirteen candidate peptides were synthesized and screened for activity against a panel of four fungal species, including both plant and human pathogens. Of the five peptides with antifungal activity, two were found to also have low cytotoxicity in a human liver cell line. Applied on a larger-scale, our computational, semi-automated approach has the potential to help fill the void in currently available antifungal countermeasures, addressing our rising vulnerability to fungal pathogens.

**Figure 1.**
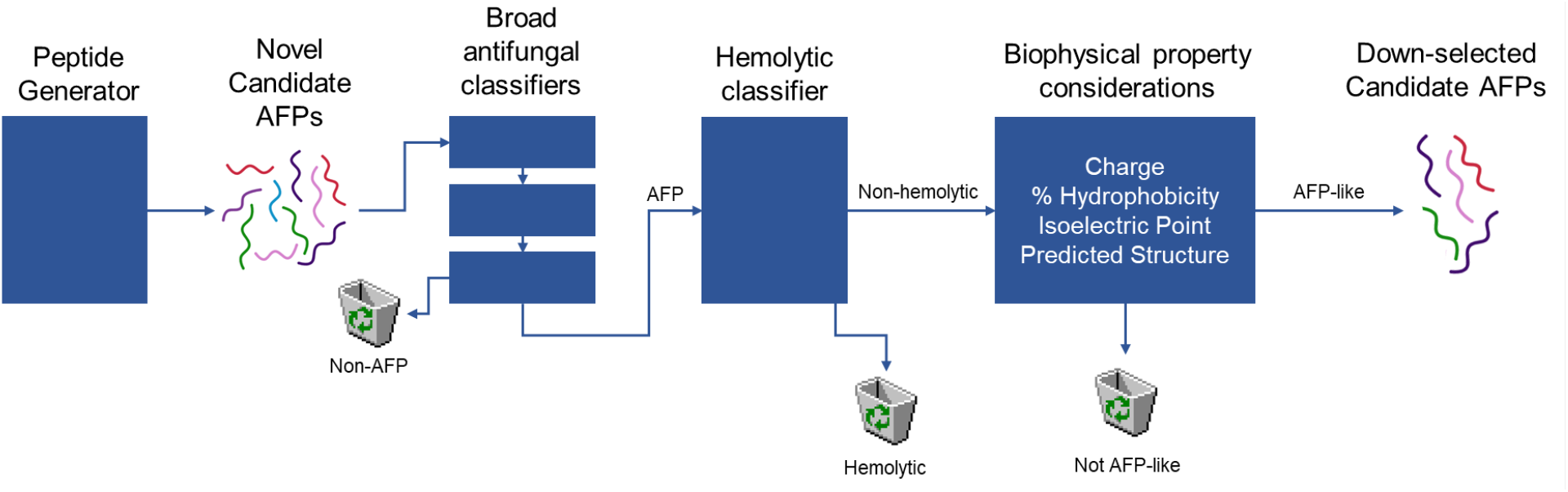
High-level overview of Fung-AI pipeline for antifungal discovery. Candidate antifungal peptides (AFPs) were generated and then down-selected by passing them through three antifungal classifiers and a hemolytic classifier. AFP-like, non-hemolytic candidates with reasonable biophysical properties and diverse secondary structures were retained for *in vitro* antifungal and cytotoxicity testing.

## Results

### In silico *generation and down-selection of candidate antifungal peptides*

A total of 9,994 unique peptide sequences ranging from 10-35 amino acids in length were computationally generated with our custom GAN (**S1 Text**). Following peptide generation, we applied a series of in-house and publicly available *in silico* tools to further screen for generated peptides that were the most likely to be antifungal, non-hemolytic (*e*.*g*., not obviously toxic to humans), and physiologically realistic. In addition, the novelty of promising candidates was assessed by querying generated peptides against known proteins in the NCBI Non-Redundant (nr) Protein Database. Cumulatively, in the first phase of this effort, the Fung-AI pipeline (**Figure 1**), was used to down-select five candidate peptides for experimental validation. More specifically, generated peptides were sequentially removed and assessed throughout the Fung-AI pipeline (**Figure 2A**). First, all 9,994 peptides were passed through three binary classifiers, which were trained to predict whether an individual peptide would have antifungal activity (**Figure 2B-D**). Using a probability cutoff of 0.5 for each model, a total of 3,578 peptides were predicted to be antifungal by all three classifiers. Notably, the probability distributions of each classifier were concentrated at the tails, indicating strong predictions for the majority of the generated peptides, as opposed to weak predictions around the classification cutoff.

**Figure 2.**
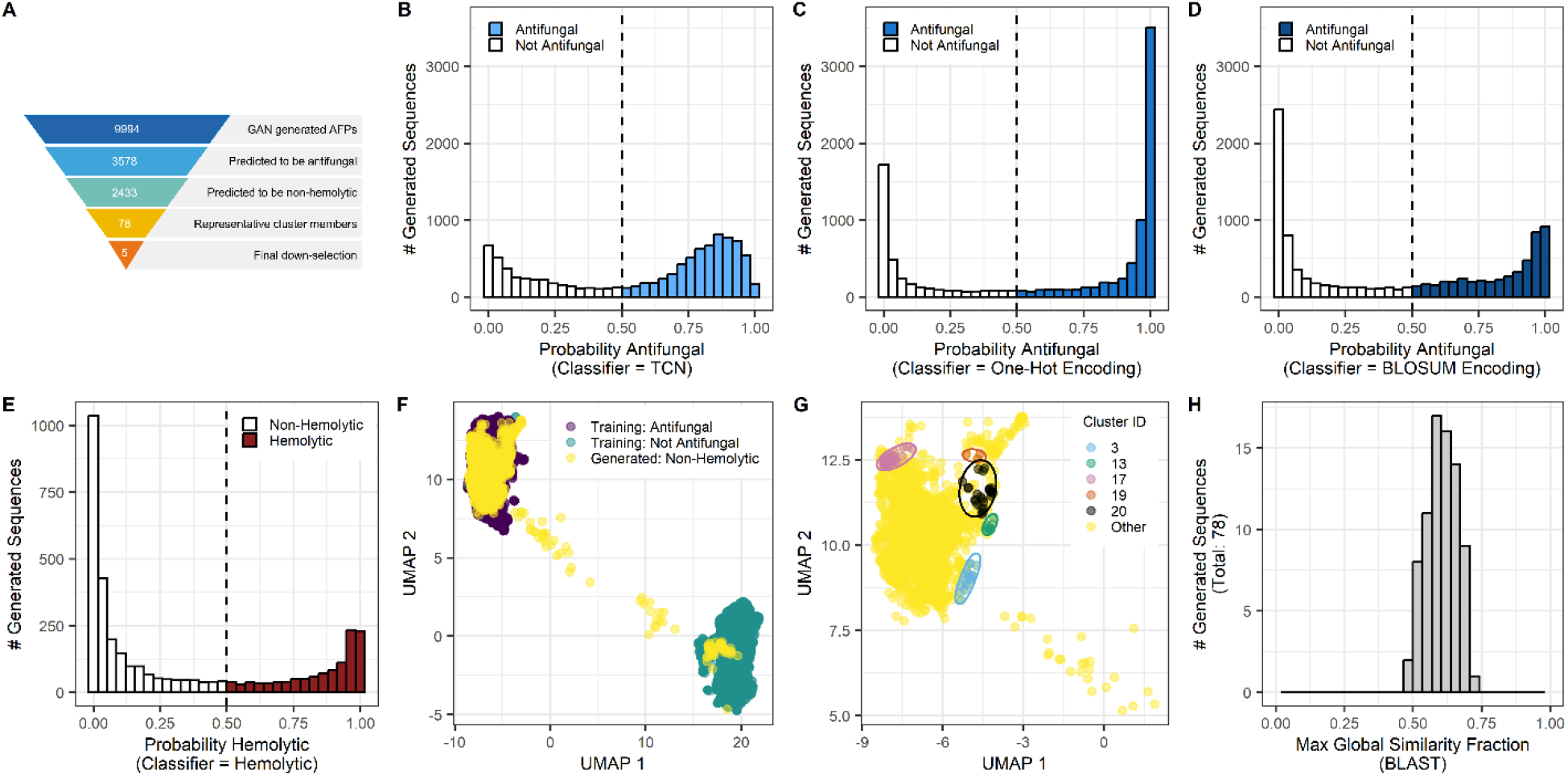
Down-selection of AI-generated candidate peptides through the *in silico* Fung-AI pipeline. **A)** Overview of the process employed to down-select candidate peptides for experimental testing. Probability distributions of 9,994 peptides being antifungal as predicted by the **B)** TCN-based, **C)** one-hot encoding-based, and **D)** BLOSUM encoding based binary antifungal classifiers. **E)** Probability distributions of 3,578 peptides being hemolytic as predicted by the binary hemolytic classifier. Classifier cutoffs were set at 0.5, denoted by vertical dashed lines. **F)** UMAP dimensionality reduction and visualization of training data compared to predicted antifungal and non-hemolytic AI-generated peptides. **G)** UMAP of five down-selected HDBSCAN clusters, containing 78 peptides of interest. Yellow denotes the rest of the generated peptides that were “AFP-like”. **H)** A total of 78 peptides representative of clusters of interest were run through BLAST. The max global similarity fraction denotes the “best” match for each generated sequence from the NCBI Non-Redundant (nr) Protein Database. The maximum possible score is 1 and the minimum possible score is 0.

A common challenge when developing antimicrobial peptides is that peptides active against microorganisms, especially fungi which are eukaryotic and thus share many characteristics with humans, may also cause cytotoxicity in mammalian cells (12). Therefore, in an attempt to increase the chances of identifying relatively safe therapeutic candidates, the remaining 3,578 peptides that were predicted to be antifungal by all three classifiers were passed through a binary hemolytic classifier to predict whether or not peptides would have hemolytic activity (**Figure 2E**). Hemolytic peptides were removed, and embeddings of the remaining 2,433 generated peptides were compared with training data using Uniform Manifold Approximation and Projection (UMAP) (**Figure 2F**). The majority of predicted antifungal and non-hemolytic generated peptides clustered with known antifungal peptides from the training data.

To identify groups of similar peptides that could be sampled for experimental validation, we used hierarchical density-based spatial clustering of applications with noise (HDBSCAN) to identify putative antifungal, non-hemolytic peptide clusters. Training data clusters of interest were defined as those which had greater than 30 antifungal member sequences, of which at least 70% were predicted to be non-hemolytic. Of the training data, nine clusters met these criteria. Generated sequences were then mapped to these clusters, with a total of 78 generated sequences mapping to five out of the nine clusters, and none mapping to the remaining four clusters (**Figure 2G**). When queried against short known proteins (< 250 amino acids), no considerable similarities (> 90% query cover and > 80% percent identity) were found between the top BLAST hits and these 78 sequences, with all max global similarity fractions below 0.75 for hits with at least 50% query cover. This suggests that our AI-generated sequences are truly novel and not simply copies or near-relatives of known natural or synthetic proteins (**Figure 2H**).

Finally, having down-selected to five candidate clusters of interest, one peptide from each cluster was selected based on a combination of biophysical properties, predicted structure, and synthesizability. Following published literature on synthetic therapeutic peptide design (11), we filtered for mildly cationic peptides (*e*.*g*., positive net charge below +6) composed of 30-60% amino acids with hydrophobic side chains (**Table 1**). The PEP-FOLD4 server (41) was used to predict the structures of the remaining peptides. For each cluster, one peptide with a qualitatively representative linear structure (*e*.*g*., alpha-helical, random coil, *etc*.) was selected for synthesis (**Figure 3**). Thus altogether, the Fung-AI pipeline is a semi-automated, fully computational approach to generating and priority ranking novel antifungal peptide candidates for experimental testing.

**Table 1.**
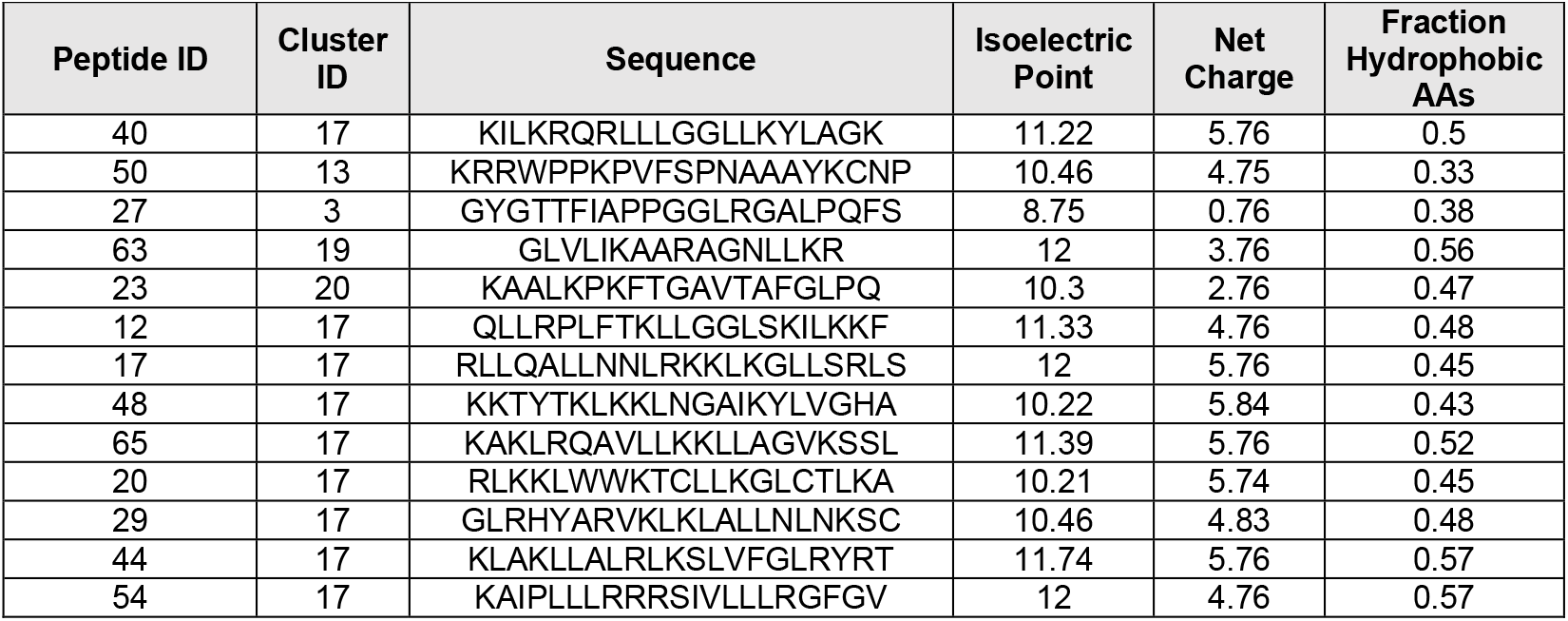
Biophysical properties of peptides selected for experimental validation. AA = amino acid.

**Figure 3.**
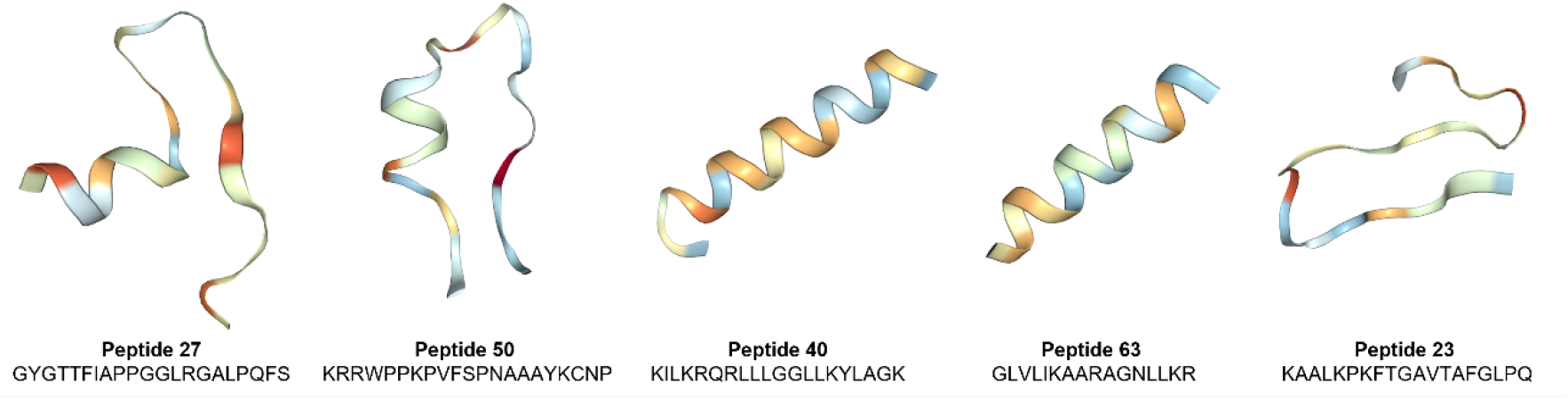
Predicted structures of down-selected AI-generated peptides. Structures predicted with the PEP-FOLD4 server. Residues are colored by hydrophobicity (red = hydrophobic, blue = hydrophilic).

### Antifungal activity and cytotoxicity of AI-generated peptides

To evaluate the antifungal potential of candidate peptides, we first measured MICs of our five down-selected AI-generated peptides against the relevant wheat pathogen, *Fusarium graminearum*, and the model organism *Saccharomyces cerevisiae*. Peptide 40 from cluster 17 showed inhibitory activity against both organisms, prompting a second phase of testing, during which time eight additional peptides from cluster 17 which met down-selection criteria were also synthesized and tested (**Figure S1**). A total of five out of nine peptides from cluster 17 showed inhibitory activity against *F. graminearum*. These five peptides were then further tested against an expanded fungal panel of human-relevant pathogens: *Candida albicans* and *Candida auris*. We finally assessed cytotoxicity of the down-selected peptides by determining the LC_50_values in HepG2 human liver carcinoma cells, where the LC_50_is defined as the lethal concentration that kills 50% of cells Peptides 12 and 40 exhibited the broadest and strongest antifungal activity with MICs of 250 µg/mL against *F. graminearum* and 500 µg/mL against both *S. cerevisiae* and *C. albicans* (**Table 2**). Peptide 48 showed weaker, but equally broad activity, inhibiting *F. graminearum, S. cerevisiae*, and *C. albicans* at 500 µg/mL. In contrast, Peptide 17 displayed activity only against *F. graminearum* at 500 µg/mL. Peptide 65 was active against *F. graminearum* and *C. albicans* at 500 µg/mL but had no effect on *S. cerevisiae*. Notably, none of the peptides inhibited *C. auris* at concentrations up to 500 µg/mL (**Figure S2**).

**Figure S1.**
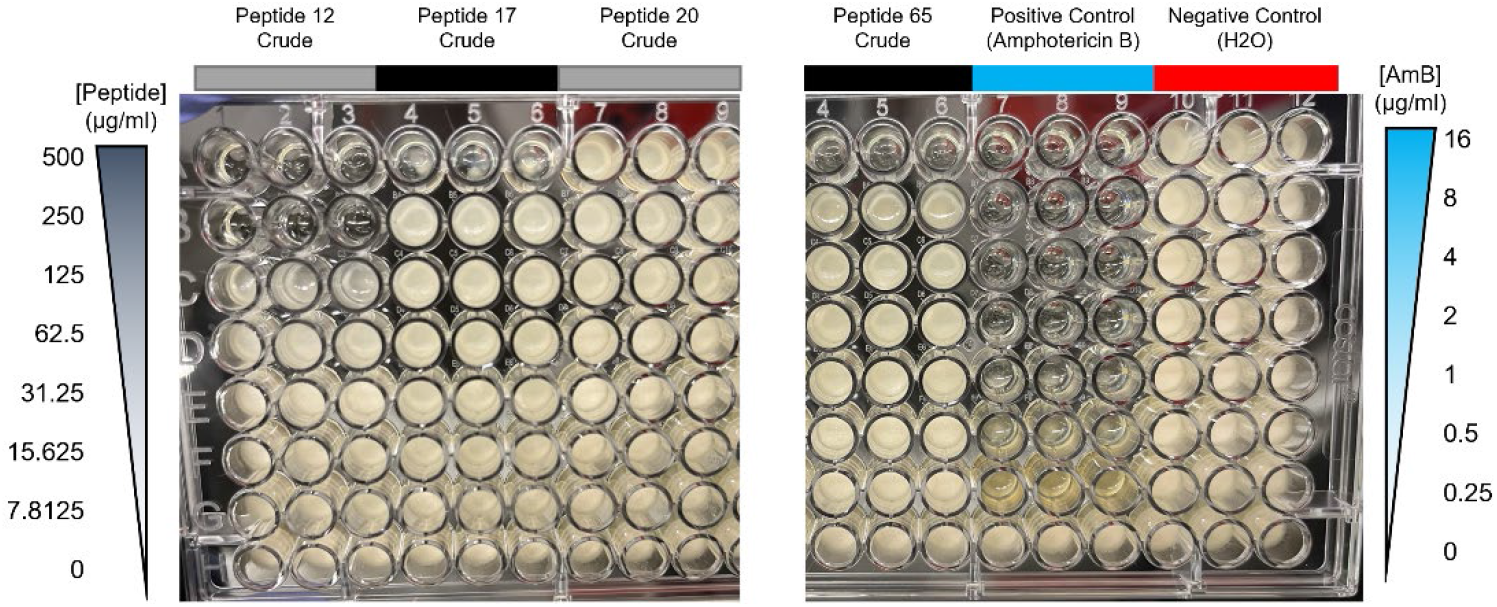
Example of the *Fusarium graminearum* MIC assay for a representative selection of active and inactive peptides. Images of the MIC assay for four peptides. Amphotericin B (AmB) was used as a positive control (blue) and water was used as a negative control. The bottom row of each plate was a no-treatment control. The MIC was reported as the lowest concentration of antifungal that inhibited growth.

**Table 2.**
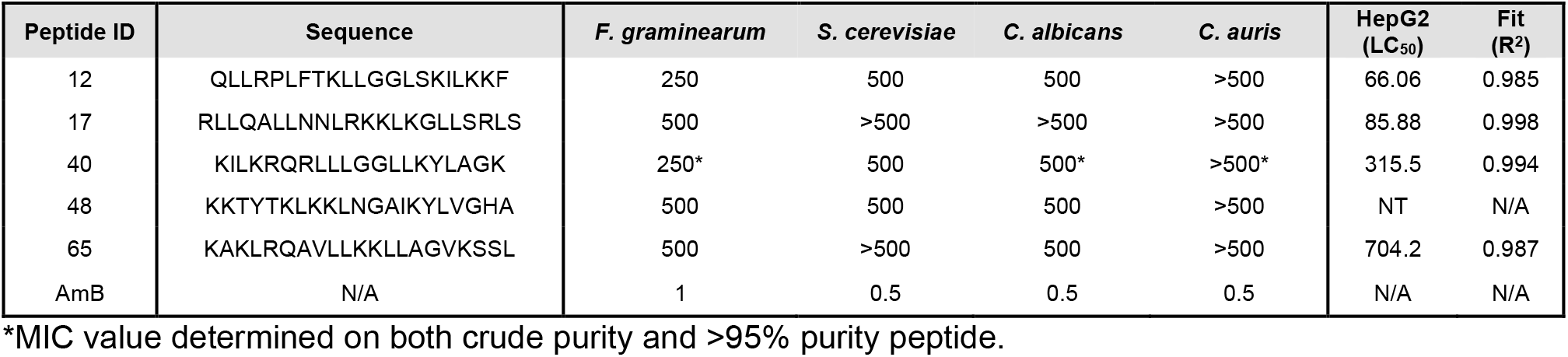
MIC (µg/mL) and LC_50_(µg/mL) activity against a panel of fungi and HepG2 cells, respectively. R^2^ denotes the fit of the dose-response curve used to calculate LC_50_values. NT: No toxicity observed. AmB: Amphotericin B. N/A: Not available.

**Figure S2.**
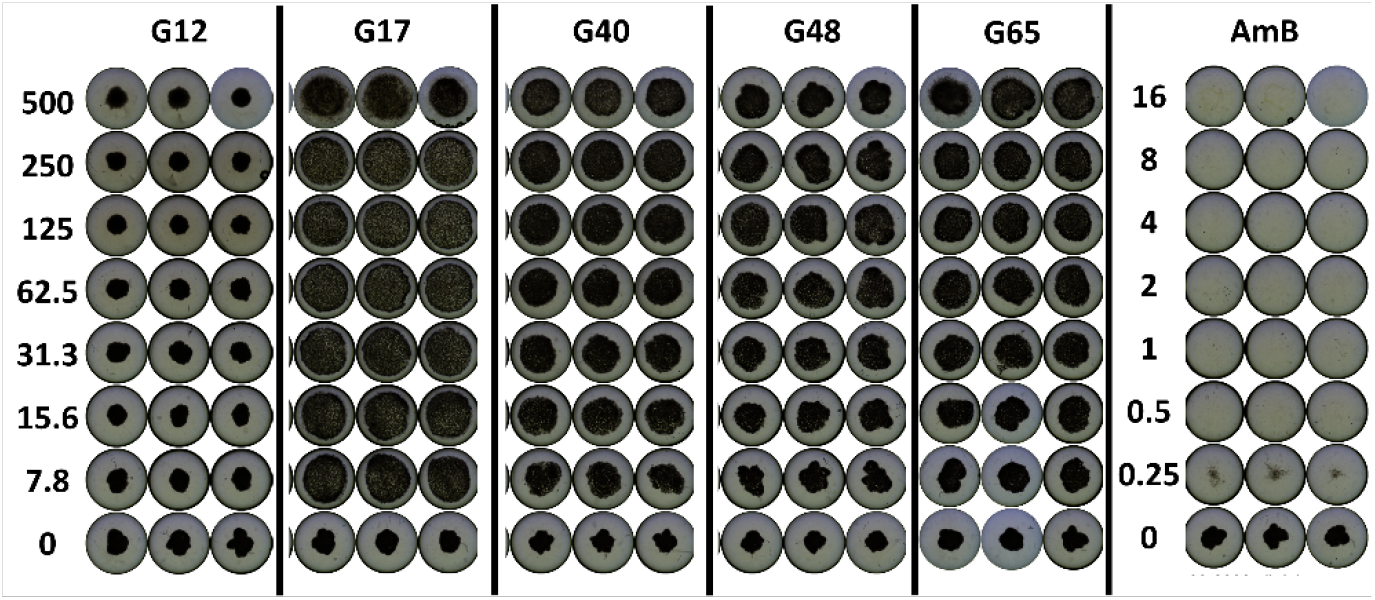
MIC assay of peptides against *Candida auris* strain CDC 381. *C. auris* is an emerging pathogen that is difficult to treat clinically. We tested peptides that were active against *F. graminearum* against *C. auris*. Amphotericin B (AmB) was used as a positive control. The bottom row of each plate was a no-treatment control. None of the tested peptides were found to inhibit this strain of *C. auris* or a more resistant isolate *C. auris* strain 390 (data not shown).

Cytotoxicity varied substantially among the peptides (**Table 2, Figure 4**). Peptide 12 had the lowest LC_50_ value (66.06 µg/mL), indicating higher cytotoxic potential in HepG2 cells. In contrast, no toxicity was observed for Peptide 48 at the concentrations tested, and Peptide 65 showed only mild cytotoxicity with an LC_50_ of 704.2 µg/mL. Peptide 40 and Peptide 17 exhibited intermediate cytotoxicity with LC_50_ values of 315.5 µg/mL and 85.88 µg/mL, respectively.

**Figure 4.**
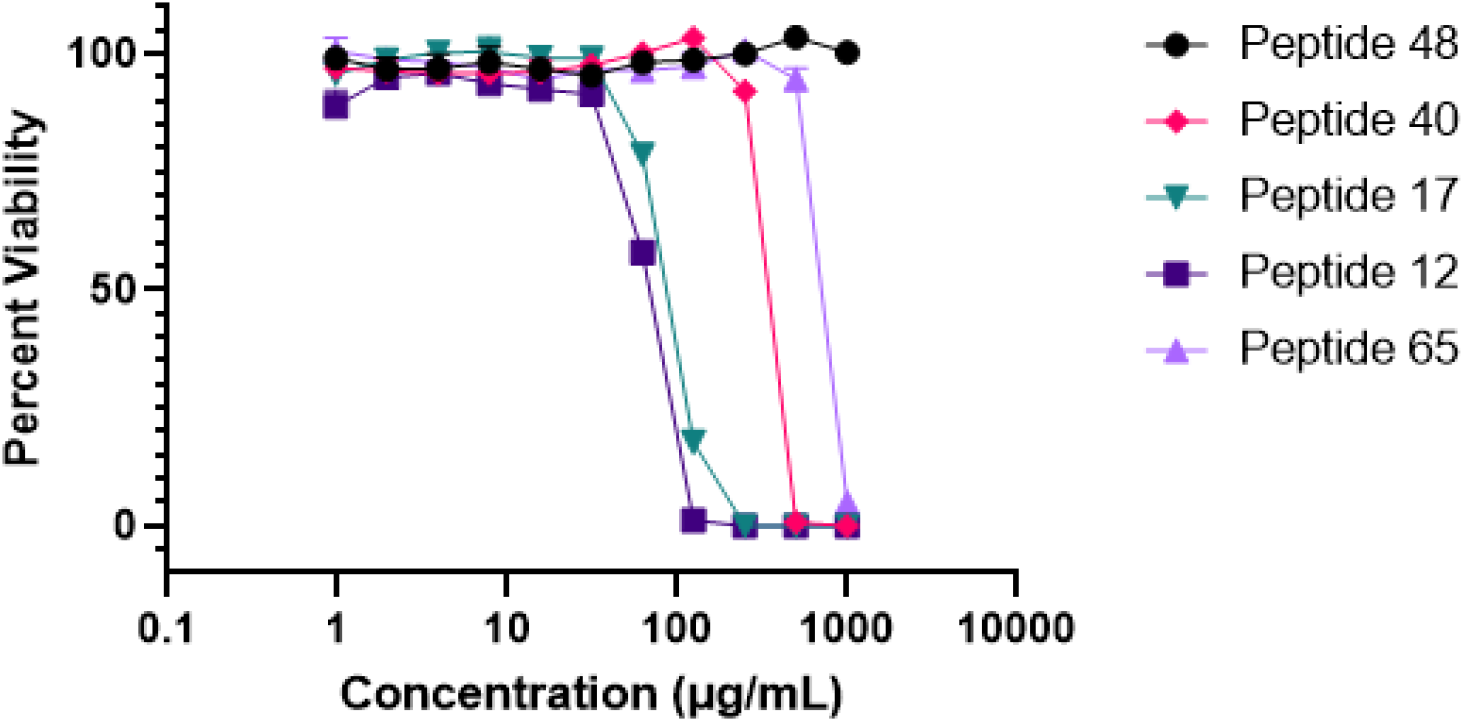
Cytotoxicity of AI-generated peptides in human liver cells. Percent viability of HepG2 cells at increasing concentrations of peptide. Each point denotes the mean and error bars denote the standard deviation across three replicate wells.

Overall, peptides 12 and 40 demonstrated modest antifungal activity, particularly against the wheat pathogen, *F. graminearum*. While these peptides were also active against *C. albicans*, their therapeutic window against human pathogens may be limited by cytotoxicity. Peptides 48 and 65, despite showing weaker antifungal activity and being predicted to form cationic alpha-helical structures (**Figure 5**), may be safer scaffolds for future optimization due to their low cytotoxicity profile.

**Figure 5.**
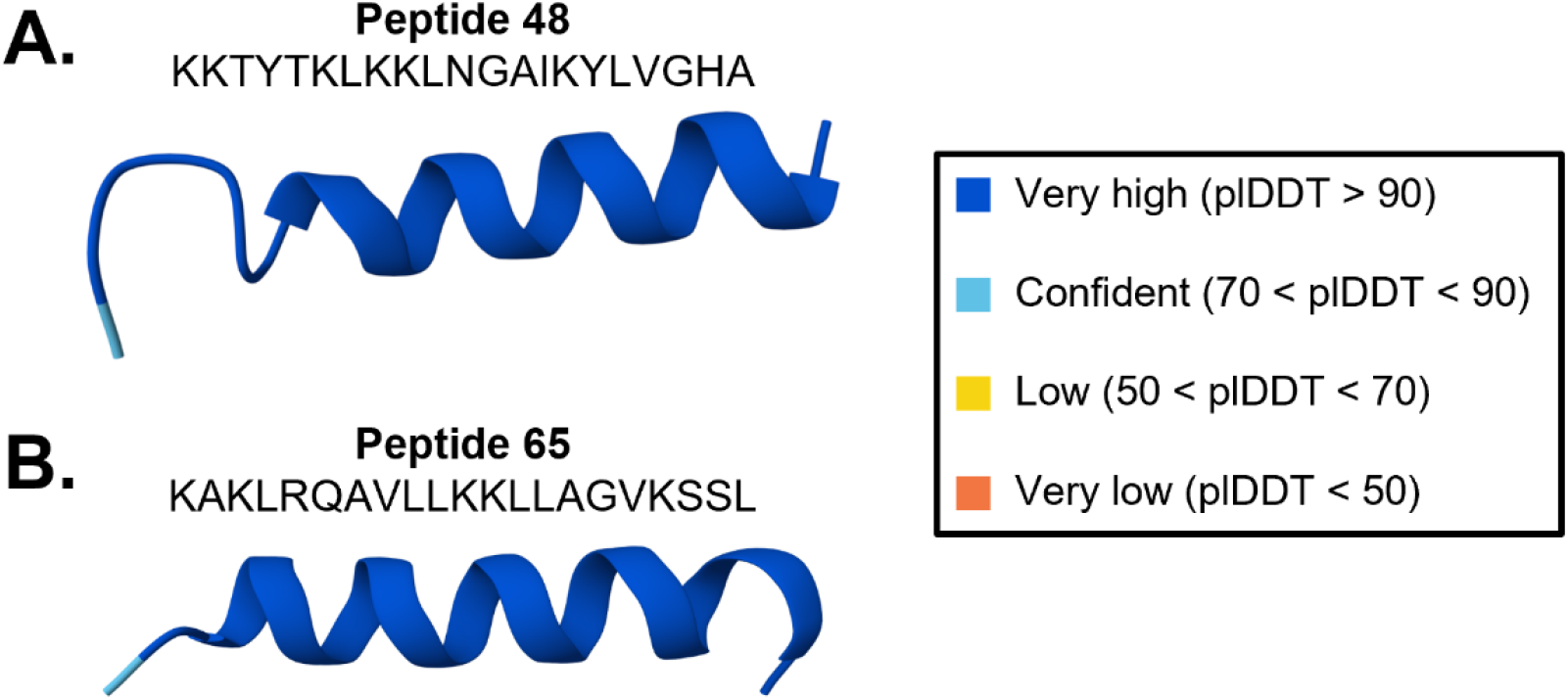
Predicted structures of antifungal and non-cytotoxic AI-generated peptides. Structures were predicted for (**A**) peptide 48 and (**B**) peptide 65 with the publicly available AlphaFold3 server. Residues are colored by the predicted local distance difference test (plDDT) confidence score. Dark blue: very high confidence (plDDT > 90). Light blue: confident (70 < plDDT < 90).

## Discussion

Here, we have developed and experimentally validated the Fung-AI pipeline, an AI/ML supported approach to design *de novo* antifungal peptides. We generated ∼10,000 candidate peptides *in silico* with a custom GAN, computationally down-selected hits, and evaluated peptide antifungal activity across a panel of fungal pathogens of relevance in agriculture and human health. Testing fewer than 20 peptides across two phases of experimental validation, we identified one cluster of peptides from which five out of nine synthesized peptides (55%) displayed activity against at least one fungal species. Of these five peptides, two displayed both antifungal activity in *F. graminearum* and *C. albicans* as well as low cytotoxicity in a human liver cell line (*e*.*g*., LC_50_ was less than the MIC), suggesting there may be the potential to further optimize these peptides for therapeutic applications (**Table 2**). Notably, while the sequences of these peptides are distinct from known proteins, they are cationic and predicted to form alpha-helical structures, meaning that they may act in a functionally similar way to known antimicrobial peptides. Overall, we have demonstrated the utility of our peptide generator and the Fung-AI pipeline by discovering novel antifungal peptides with minimal experimental screening.

Through building the Fung-AI pipeline, we have identified three key limitations of our approach as well as opportunities to improve the usefulness of AI-guided approaches for antifungal discovery. First, publicly available antifungal peptide data is limited. For example, while there are over ∼35,000 antimicrobial peptides in the data repository of antimicrobial peptides (DRAMP) database, only ∼5% of the peptides in DRAMP are labeled as antifungal (42). Therefore, in order to maximize the size of our training dataset, our GAN was trained on any known antifungal peptides, regardless of target species. While it is interesting that such a heterogenous training set resulted in the generation of peptides that were broadly active against both *F. graminearum* and *C. albicans*, our peptides had relatively high MICs (**Table 2**) compared to known antifungal peptides such as VLL-28 (MIC *C. albicans*: 88.5 µg/mL) (13). Using smaller pathogen-specific data to fine-tune our GAN or develop additional antifungal classifiers could potentially increase the chances of generating targeted, high-potency therapeutics. In addition, it may be possible to modify these peptides to increase their activity. Alternatively, a recent study by Wang *et al*. found that training a diffusion model on a broad collection of antimicrobial peptides (antibacterial and antifungal) was sufficient to discover peptides with activity against *C. glabrata* and *Cryptococcus neoformans* (43).

Second, while based on their broad-spectrum activity, positive charge, and predicted linear alpha-helical structures, we hypothesize that peptides 48 and 65 may be membrane disrupting, we did not experimentally determine the mechanism of action (MOA) of any of our AI-generated peptides. Those looking to leverage our pipeline or similar methods should aim to incorporate MOA prediction into the down-selection process. Additionally, newer generative-AI methods such as protein diffusion models (17) or small molecule diffusion models (44,45) could be explored to generate peptides or small molecule drugs against specific fungal targets.

Finally, none of the peptides evaluated in this work were found to considerably inhibit the growth of *C. auris*, a fungal pathogen of growing clinical concern that often shows a multi-drug resistance phenotype. *C. auris* was first described in 2009 (46) and consequently, publicly available data sets on this pathogen are fairly scarce. For example, only 78 peptides out of ∼24,000 total entries contained in the Database of Antimicrobial Activity and Structure of Peptides (DBAASP) (47) are associated with any strain of *C. auris*. Future studies should aim to collect high-throughput, high-quality multi-modal data (*e*.*g*., small molecule antifungal susceptibility, antifungal peptide screens, whole-genome sequencing data, clinical metadata, *etc*.) for this pathogen to enable the development of targeted and effective therapeutic strategies.

In summary, the Fung-AI pipeline demonstrates the value of applying generative AI toward the challenge of antifungal drug discovery. Our results motivate the continued exploration, screening, and optimization of AI-generated peptides against relevant agricultural and human fungal pathogens.

## Materials and Methods

### Construction of the in silico Fung-AI pipeline

#### Dataset curation

Two datasets were created for this work: one capturing antifungal peptides and non-antifungal peptides, and another of hemolytic peptides and non-hemolytic peptides.

The antifungal peptide dataset consisted of 9,388 peptides from published literature (35), DRAMP 4.0 (42), and CAMPR4 (48). Antifungal peptides from CAMPR4 and DRAMP 4.0 could be naturally occurring or synthetic, but had to have experimentally validated antifungal properties. Peptides only hypothesized to be antifungal were not included in the dataset. A total of 2,204 peptides were gathered from the CAMPR4 database, of which 982 (44.6%) were antifungal and 1,222 (55.4%) were non-antifungal. An additional 1,811 peptides, of which all were antifungal, were pulled from the DRAMP 4.0 database. Finally, 5,373 peptides from Sharma et al. (37) were included, of which 1,932 (36.0%) were antifungal and 3,441(64.0%) were non-antifungal. In all, the peptides ranged from 2 to 726 amino acids in length, with a mean length of 26.0 (σ = 31.5) and a median length of 20.0 amino acids. As this distribution is heavily skewed with a very long tail, we set a minimum peptide length of 10 and a maximum of 35 amino acids. After removing sequences that were too long or too short, the final dataset contained 7,335 peptides, of which 3,423 were antifungal peptides (46.7%) and 3,912 were non-antifungal peptides (53.3%). The data was split into training and test sets using an 80/20 split.

A dataset of hemolytic and non-hemolytic peptides published by Yaseen *et al*. (49) was used in the development of our hemolytic classifier. This dataset contained 3,804 peptides, of which 1,576 (41.4%) were hemolytic and 2,228 (58.6%) were non-hemolytic. The mean peptide sequence length was 18.3 (σ = 6.2) amino acids with a maximum length of 35 and a minimum of 10 amino acids. The data was split into training and test sets using an 80/20 split.

#### Antifungal peptide classifiers

Three antifungal peptide classifiers were identified from literature, retrained using our antifungal peptide dataset, and implemented within the Fung-AI pipeline. All three classifiers were used to down-select AI-generated candidate peptides, while one model was additionally used to monitor the behavior of the GAN throughout training. Detailed descriptions of the architectures and training regimens for each model are provided in the **Supporting Information (S1 Text)**.

The first model, based on temporal convolutional networks (TCNs) (50) leveraged an architecture described by Singh et al (36). Following retraining, the TCN model achieved an accuracy of 86.8%, area under the curve (AUC) of 92.5%, and F1-score of 86.1% on our antifungal peptide datasets.

The second and third models used the architecture described in Sharma et al. (37), which is based on a one-dimensional convolutional neural network bidirectional long short-term memory (1DCNN-BiLSTM) architecture. The two models were differentiated by the input and the presence of an embedding layer. The second model took as input the one-hot encoded versions of the peptides, forgoing the need for an embedding layer. The third model used an embedding layer with a dimension of 20 and was pre-seeded with the BLOSUM weights (51). On our antifungal peptide dataset, the second and third models achieved accuracies of 85.5% and 86.3%, AUCs of 92.1% and 92.4%, and F1-scores of 84.6% and 84.7% respectively.

#### Hemolytic classifier

The hemolytic peptide classifier is based on published work by Yaseen et al. (49), but modified to use the architecture of the third antifungal peptide classifier (1DCNN-BiLSTM, using the input based on BLOSUM weights) (51). We were able to achieve comparable results to those referenced in the paper using this architecture. The model was trained on a random subset of 80% of the hemolytic peptide dataset for 20 epochs with a batch size of 16, using the Adam optimizer with a learning rate of 0.03. The remaining 20% of the data was used for testing. The model achieved an accuracy of 77.9%, AUC of 85.9%, and F1-score of 78.3%, as compared to the 88% AUC reported in Yaseen et al. (49).

#### Generative adversarial network

A GAN is a generative model framework composed of two neural networks, a generator *G*, and a discriminator *D*. The two models are trained alternatingly, such that they are in a zero-sum game in which the generator tries to create realistic looking outputs and the discriminator tries to identify real and fake data. The generator creates fake data, the discriminator is trained on real and fake data, then the discriminator weights are frozen and the generator is trained to fool the discriminator. This process repeats until the model reaches a point where the loss function is not improving with additional training.

Unlike standard GANs, our model has an additional encoder that pairs with the generator, which in turn acts as both a generator and a decoder. As the dataset was somewhat small, which resulted in poor performance when trained solely on antifungal peptides, we applied two strategies to create a larger dataset for training the GAN. First, we trained on both antifungal and non-antifungal peptides, accounting for this in the discriminator loss function. Second, we trained an autoencoder while simultaneously training the generator, which had the benefit of reducing mode collapse. The model was forced to still be able to reconstruct real antifungal and non-antifungal peptides, thereby enforcing meaning in the embedding space. As a result, there are three models that compose the GAN: the discriminator, the generator/decoder, and the encoder. This setup prevented mode collapse and enabled the generation of diverse output sequences. Detailed descriptions of the GAN architecture, loss function, and training methods are provided in **S1 Text**.

#### Unsupervised clustering of peptide sequences

The BLOSUM embedding layer from the 1DCNN-BiLSTM antifungal classifier was used to generate embeddings for both the known and GAN-output peptide sequences. UMAP was then used to reduce the peptide embeddings down to two dimensions, fitting the UMAP model to classifier training data (52). Following feature compression with UMAP, the hierarchical density-based spatial clustering of applications with noise (HDBSCAN) algorithm was used to cluster the training data with the following parameter settings: min_cluster_size = 40, min_samples = 2, gen_min_span_tree = True, and prediction_data = True. Generated peptides were then assigned to the existing clusters for visualization and down-selection for experimental testing. HDBSCAN version 0.8.33 (53), UMAP-learn version 0.5.5 (52) were implemented in Python version 3.9.13.

#### Peptide alignment to known proteins

In order to characterize the relative novelty of the AI-generated peptides, the Basic Local Alignment Search Tool (BLAST) version 2.15.0 (54) was used to compare generated peptides to documented protein sequences found in nature. A maximum of 100 BLAST sequences were returned for each query, and reference hits greater than 250 amino acids or with less than 50% query cover were removed.

An additional metric was developed to quantitively determine the “best” match for each generated sequence when queried against the standard NCBI Non-Redundant (nr) Protein Database used by BLAST. This metric, referred to here as the “Global Similarity Fraction”, is the scaled product of two BLAST alignment metrics: query cover and percent identify:

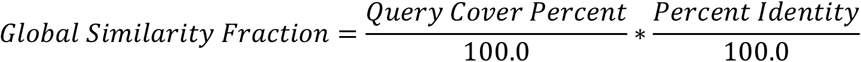

where the query cover is defined as the percent of the generated sequence that was found in the returned BLAST sequence and the percent identity is defined as the percent of the returned BLAST sequence that identically matches the generated sequence. For each generated sequence, the BLAST result with the maximum Global Similarity Fraction was assigned as the “best”, or most similar, reference protein sequence.

#### Biophysical property calculations

Biophysical properties including net charge, isoelectric point, and the percent of hydrophobic residues were calculated for a subset of peptide sequences during down-selection for experimental validation. The percent of hydrophobic residues was calculated from peptide sequence, based on the number of amino acids containing aliphatic or aromatic hydrophobic side chains (A, I, L, M, V, F, W, and Y) (55). Net charges and isoelectric points were calculated with the Biopython package version 1.83 (56). Positively charged (*e*.*g*., 0 < net charge < +6) peptides composed of 30-60% hydrophobic amino acids were further considered for experimental testing.

#### Prediction of peptide structures

The publicly available server-based tool for peptide structural prediction, PEPFOLD-4 (41) was used to predict the structures of candidate peptides of interest during initial down-selection for experimental testing. The AlphaFold3 server (57) was used to predict and visualize the structures of peptides following experimental validation.

### Experimental validation of the Fung-AI pipeline

#### Fungal strains

*Saccharomyces cerevisiae* S288C (ATCC; 204508), *Candida albicans* 23Q (BEI Resources; NR-29341) and *Candida auris* CDC 381 (Casadevall Lab, CDC & FDA Antibiotic Resistance Isolate Bank) were cultured with Yeast Peptone Dextrose (YPD) media at 30°C. *Fusarium graminearum* PH-1 (ATCC; MYA-4620) was maintained on Potato Dextrose Agar (PDA) at 30°C.

#### Peptide synthesis preparation

The tested peptides in **Table 3** were synthesized (ABI Scientific) with either crude purity and desalting for the initial assays, or with 95% purity for hit validation assays. A 10 mg/mL stock solution of each peptide was prepared in filter sterilized Milli-Q water. Stock solutions were subsequently diluted in YPD media to obtain 1 mg/mL working solutions.

**Table 3.**
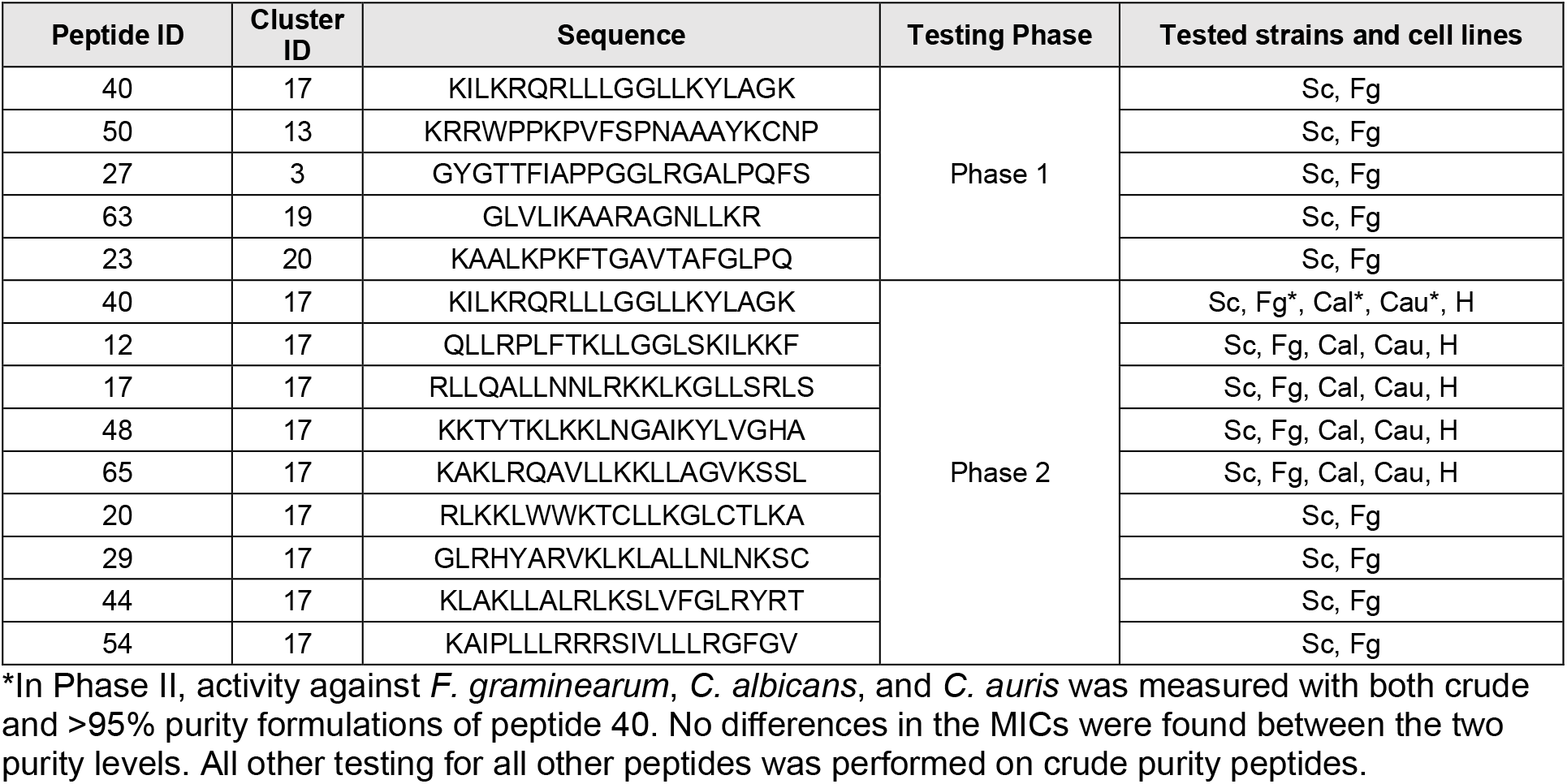
Selection of peptides tested for inhibitory activity and/or cytotoxicity. Testing phase denotes whether peptides were from original screen or deep dive into promising candidate clusters. Sc = *Saccharomyces cerevisiae*, Fg = *Fusarium graminearum*, Cal= *Candida albicans*, Cau = *Candida auris*, H = HepG2 cells.

#### Minimum inhibitory concentration (MIC) assays

Inoculums for *S. cerevisiae, C. albicans*, and *C. auris* were prepared by diluting an overnight culture 1:100 in YPD media and incubating until cultures reached an OD600 of 0.2. MIC assays for *F. graminearum* PH-1 were adapted from Al-Hatmi, et al (58). Briefly, *F. graminearum* PH-1 spores were prepared by streaking *Fusarium* on a PDA plate and incubating at room temperature for 5 days. Spores were scraped into a solution of Phosphate Buffered Saline (PBS) with 0.005% Tween 20. The solution was allowed to settle for three minutes. The upper solution was carefully transferred to a new tube and vortexed for 15 seconds. The spore suspension was adjusted to an OD530 of 0.15 in YPD media.

For *S. cerevisiae, C. albicans*, and *F. graminearum*, 100 µL of the diluted fungal culture or spore solution was added to each well of a microtiter plate. A 100 µL aliquot of the peptide working solution (1 mg/mL) or amphotericin B positive control (32 µg/mL) was added to the first row, with each condition tested in triplicate. For *C. auris*, 50 µL of diluted fungal culture and peptide working solution were added to the wells for a total assay volume of 100 µL. A two-fold serial dilution was performed down to the 7^th^ row, leaving the last row as a no-treatment control, giving a tested range of 500-7.8 µg/mL for the peptides and 16-0.25 µg/mL for amphotericin B. *S. cerevisiae, C. albicans* and *C. auris* microtiter plates were incubated at 28°C for 48 hours and visually inspected for growth, while *F. graminearum* was incubated at 34°C. MIC is reported as the lowest concentration of antifungal that inhibited growth.

#### Cell cytotoxicity assays

To assess the toxicity of peptides in mammalian cell culture, a cytotoxicity assay was performed. Briefly, HepG2 cells (ATCC; HB-8065) were seeded in a 96-well plate at a density of 25,000 cells per well and allowed to grow overnight at 37 °C, 5% CO_2_. Eagle’s minimum Essential Medium (EMEM) with 10% fetal bovine serum (FBS) was used for cell growth. Peptides of interest were diluted serially from 1000 to 0.98 μg/mL with a twofold dilution scheme. For testing, 100 µL of the diluted peptide solution was added to each well, with each concentration tested in triplicate. Each assay plate included the appropriate negative, positive and vehicle controls. The samples and cells were allowed to incubate at for 24 hours at 37°C prior to a viability assessment with CellTiter Blue (Promega). CellTiter Blue was used in accordance with manufacturer’s recommendations with a four-hour incubation time.

Percent viability was calculated by comparing the fluorescence value of the test well to that of the negative control. Each concentration was tested in triplicate (n = 3 wells). Dose-response curves were fit on averaged viability data for each peptide using a pre-defined non-linear regression model in Prism (Model: [Inhibitor] vs. normalized response – Variable slope). The LD_50_ (µg/mL) and model fit (R^2^) were calculated from each model.

## Data Availability

All data used in model development and figure generation can be found at https://github.com/jhuapl-bio/fungai.

## Code Availability

All code used to train models and generate figures is publicly available at https://github.com/jhuapl-bio/fungai.

## Supporting Information

*S1 Text:* Document providing detailed descriptions of AI-based methods used in the Fung-AI pipeline.

*Figure S1*: Example of the *Fusarium graminearum* MIC assay for a representative selection of active and inactive peptides.

*Figure S2*: MIC assay of peptides against *Candida auris* strain CDC 381

